# Gradual proactive regulation of body state by reinforcement learning of homeostasis

**DOI:** 10.1101/2025.10.17.682979

**Authors:** Mana Fujiwara, Honda Naoki

## Abstract

Living systems maintain physiological variables such as temperature, blood pressure, and glucose within narrow ranges—a process known as homeostasis. Homeostasis involves not only reactive feedback but also anticipatory adjustments shaped by experience. Prior homeostatic reinforcement learning (HRL) models have provided a computational account of anticipatory regulation under homeostatic challenges. However, existing formulations lack mechanisms for gradual, trial-by-trial adjustment and for extinction learning.

To address this issue, we developed a continuous HRL framework that enables trial-wise tuning of anticipatory regulation. The model incorporates biologically informed components: asymmetric reinforcement, weighting negative outcomes more than positive outcomes; and a dual-unit, context-gated inhibitory mechanism.

We applied the framework to thermoregulatory conditioning with ethanol-induced hypothermia and successfully reproduced cue-triggered compensation, gradual tolerance, and rapid reacquisition after extinction. We then extended the framework to multiple physiological variables influenced by shared neural or hormonal control signals, and found that when regulatory priorities across variables were uneven, a deviation in one variable propagated through shared control to others, yielding a cascading, system-wide failure to return to ideal state (non-recovery)—a pattern reminiscent of autonomic dysregulation (e.g., dysautonomia, ME/CFS). Overall, our framework provides a computational basis to advances a systems-level understanding of multi-organ homeostatic dysregulation in vivo.

## Introduction

Living organisms maintain internal variables such as body temperature, blood pressure, glucose levels, and hormone concentrations within narrow ranges that are optimal for survival and proper functioning—a process known as *homeostasis*. To maintain homeostasis in dynamically changing environments, organisms rely not only on *reactive* mechanisms that respond to disturbances, but also through *proactive* control that anticipates and prepares for upcoming changes. Such proactive regulation is acquired through experience and learning.

A familiar example of this process is the development of pavlovian-conditioned drug tolerance(Farahbakhsh and Siciliano, 2023; Hou et al., 2023; Le et al., 1979; Siegel, 2001). Through repeated administrations, the body learns to adjust and attenuate the drug’s effect, eventually requiring increased dosage to achieve the same therapeutic outcome. However, upon moving to another context, which has never been associated with the drug, the same dosage may cause excessive effects, because the learned tolerance was not expressed in the new context. Another illustrative example in laboratory situations is thermoregulatory classical conditioning (Mansfield et al., 1983; Mansfield and Cunningham, 1980). In this paradigm, a neutral cue (CS) is paired with an ethanol injection (unconditioned stimulus, US) that induces hypothermia, and over repeated pairings, animals begin to generate a compensatory response to the CS by increasing body temperature in advance to counteract the expected drop, eventually leading to full tolerance. These examples illustrate that homeostatic responses can be modulated through learning in a *cue-dependent* and *anticipatory* manner. Despite its biological importance, the computational mechanisms underlying such proactive, experience-driven regulation remain largely unknown.

To address such cue-dependent and anticipatory homeostatic regulation, Keramati and Gutkin proposed a computational decision-making model of homeostasis grounded in the reinforcement learning (RL) framework (Keramati and Gutkin, 2014). In their *homeostatic reinforcement learning* (*HRL*) model, the agent learns to select actions that reduce the deviation from the optimal internal state, which are considered rewarding and reinforced, while actions that increase the deviation are punishing and thus avoided. This model has been applied to various physiological domains, including water balance, thermoregulation, and pathological conditions such as cocaine addiction (Keramati et al., 2017). Building on this framework, Uchida *et al*. extended the HRL model to account for salt appetite, modeling how the conditioned stimulus (CS) of salt taste predicts future sodium intake (Uchida et al., 2022).

While influential, the original HRL formulations face three key limitations when applied to biological systems. First, they operate with discrete actions, which makes it difficult to capture the gradual, trial-by-trial adjustment of compensatory signals. To enable gradual learning, we combined a neural circuit model with the HRL framework. The compensatory signal strength driven by the CS is updated through a reinforcement learning process to maximize net reward. This architecture enables the agent to acquire anticipatory, context-dependent control of internal states and to flexibly modulate learned responses.

Second, the previous HRL framework commonly assumes symmetric valuation of positive (reward) and negative (punishment) outcomes. However, behavioral and neural studies, particularly in decision-making tasks such as gambling, have consistently shown loss-dominant asymmetry, in which losses are weighted more heavily than equivalent gains (Macpherson et al., 2014; Sokol-Hessner and Rutledge, 2019). We extend this idea to the regulation of internal physiological states. Homeostatic systems are organized to defend safe operating ranges and to prevent critical thresholds from being crossed. Such regulation is robust: for example, body temperature is tightly maintained within 35–38 °C despite external perturbations. From a control-theoretic perspective, however, if a deviation does occur, rapidly restoring the variable risks overshoot and instability. Clinically, recovery from critical illness and multi-organ dysfunction is often protracted and gradual, with persistent organ dysfunction and long-lasting impairments documented in chronic or persistent critical illness (Iwashyna, 2012; Voiriot et al., 2022). We therefore interpret this asymmetry in control—strong defense against boundary crossings versus cautious correction once deviation has occurred—as a form of loss-dominant regulation, and we incorporate this principle into our model.

Finally, the previous HRL model does not accommodate *inhibitory learning*, such as memory extinction. In classical conditioning, xtinction refers to the gradual reduction in conditioned responses when the CS is no longer followed by the US. However, the extinguished response can re-emerge as spontaneous recovery after a delay, reinstatement after unsignaled US re-exposure, and rapid reacquisition when CS– US pairings are resumed (Myers and Davis, 2002; Napier et al., 1992; Rescorla and Heth, 1975). This phenomenon suggests that extinction does not erase the original memory, but rather involves the formation of a separate inhibitory trace, which can be unmasked by contextual cues. In the mammalian brain, such dual-memory systems are supported by the direct (excitatory) and indirect (inhibitory) pathways in the basal ganglia, mediated by D1 and D2 receptor-expressing neurons, respectively. This architecture highlights the need for models that can represent *both activation and inhibition* in associative learning (Li et al., 2016) and context-sensitive gating mechanisms are likely essential for biologically plausible extinction learning.

Here, we propose an extended HRL framework that addresses these three limitations. Our model allows (1) *gradual*, trial-by-trial updating of internal control signals; (2) *asymmetric reward weighting*, with a stronger emphasis on punishment; and (3) *inhibitory learning*, implemented through a gating mechanism that suppresses conditioned responses in the absence of expected outcomes. Then, we further extend this model to multi-dimensional homeostatic systems and demonstrate that local optimization can give rise to emergent trade-offs or even maladaptive, pathological states at the system level.

## Results

### Mathematical model of proactive homeostatic regulation

To investigate how a self-regulating system actively controls internal states in anticipation of external disturbances, we developed a computational framework based on homeostatic reinforcement learning. The model’s objective is to maintain body states within a desirable range, even in the force of perturbations. Through repeated experience, the agent learns to predict upcoming disturbances, and activates a resistance function in advance to prevent dramatic state deviations.

We first focused on thermoregulation in the context of classical conditioning. In this task, a neutral auditory cue (conditioned stimulus, CS) was repeatedly paired with an ethanol injection (unconditioned stimulus, US), which consistently induced hypothermia of approximately 1.5°C (Fig. 1A). Over successive trials, animals learned to anticipate the hypothermic effect by generating a compensatory heating response at the time of the CS presentation, ultimately developing tolerance to the US (Fig. 1B).

**Fig. 1.**
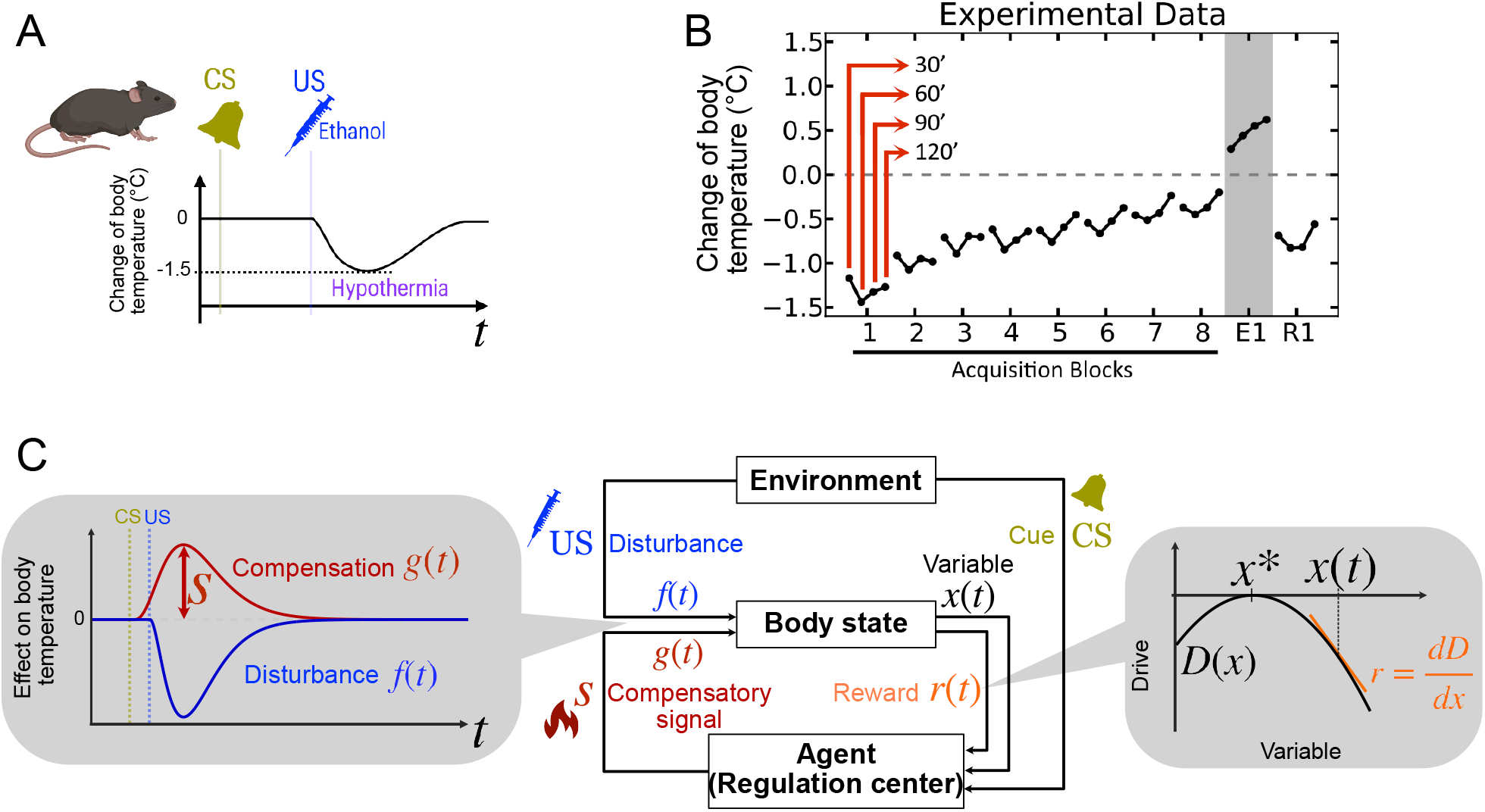
Body temperature regulation as an example of proactive homeostasis. (A) Sketch of conditioning scheme and body temperature change during the task. (B) Experimental data adapted from Karamati and Gutkin (2014), which is modified from Mansfield & Cunningham (1980). The mouse gradually obtained tolerance against ethanol-induced hypothermia, and such tolerance is included by the bell ring as shown in the extinction block (E1 after the 8th block). The 10 th trial, R1, represents the first trial of reacquisition. (C) Schematic of the proposed homeostatic RL model. Center: Block diagram illustrating interactions among the environment, body state, and agent. Left: At CS onset, a compensatory signal *g*(*t*) with the amplitude *s* is elicited, aiming to counteract the disturbance *f*(*t*) starting at US. Right: The reward *r*(*t*) is defined as the local slope of the drive function *D*(*x*) at the current state *x*(*t*) which determines the learning signal for policy adaptation.

The dynamics of body temperature *x*(*t*) in *i*-th trial is described by a differential equation

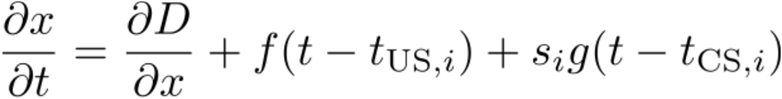

where *f*(*t*) represents the effect of the ethanol-induced disturbance and *g*(*t*) represents the compensatory response, scaled by compensatory signal *s*. The timings *t*_US,*i*_ and *t*_CS,*i*_ denote the timings of US and CS in the *i*-th trial, respectively. To represent motivational force toward the homeostatic setpoint *x*^*^, we introduced a quadratic drive function *D*+*x*(*t*)) = *a*+*x*(*t*) − *x*^*^)^2^. The compensatory signal *s*, which is generated by two opposing neural units: an activation unit *A* and an inhibition unit *I* as

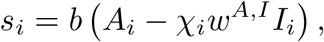

where they are up-regulated by CS, *w*^*A,I*^ represents the inhibitory effect of *I* on *A. χ* is a context-dependent gating function as *χ*_*i*_ = CS_*i*–1_US_*i*–1_, which is 1 and 0 when the previous trial was conditioning and extinction, respectively, and *b* is a constant parameter.

The agent adjust the compensatory signal *s* to maximize the cumulative of instantaneous reward, which is defined by the temporal derivative of the drive function as

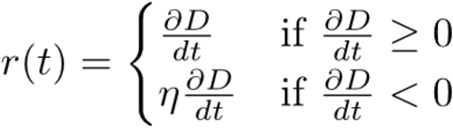

where *η* > 1 introduces an asymmetry that reflects a greater sensitivity to punishment than reward. This weighting scheme embeds a biologically motivated risk-averse strategy, whereby punishment avoidance is prioritized over reward acquisition.

To resolve the ambiguity between self-induced and externally induced changes, we introduced a context-sensitive learning rule that modulates the update of the compensatory signal *s*, depending on the inferred attribution of outcome shifts. This mechanism prevents inappropriate reinforcement when deviations arise from external disturbances rather than internal regulation (see Methods for details).

### Simulations of Proactive Body Temperature Regulation

We validated the proposed HRL model by simulating thermoregulatory learning in mice, examining whether the model can reproduce gradual acquisition of tolerance against hypothermia. In repeated CS-US pairings, the simulated body temperature showed progressive tolerance: the magnitude of the hypothermic deviation decreased across trials as the agent learned to proactively activate the compensatory response (Fig. 2A, trial 1-10), replicating empirical data (Fig. 1B).

**Fig. 2:**
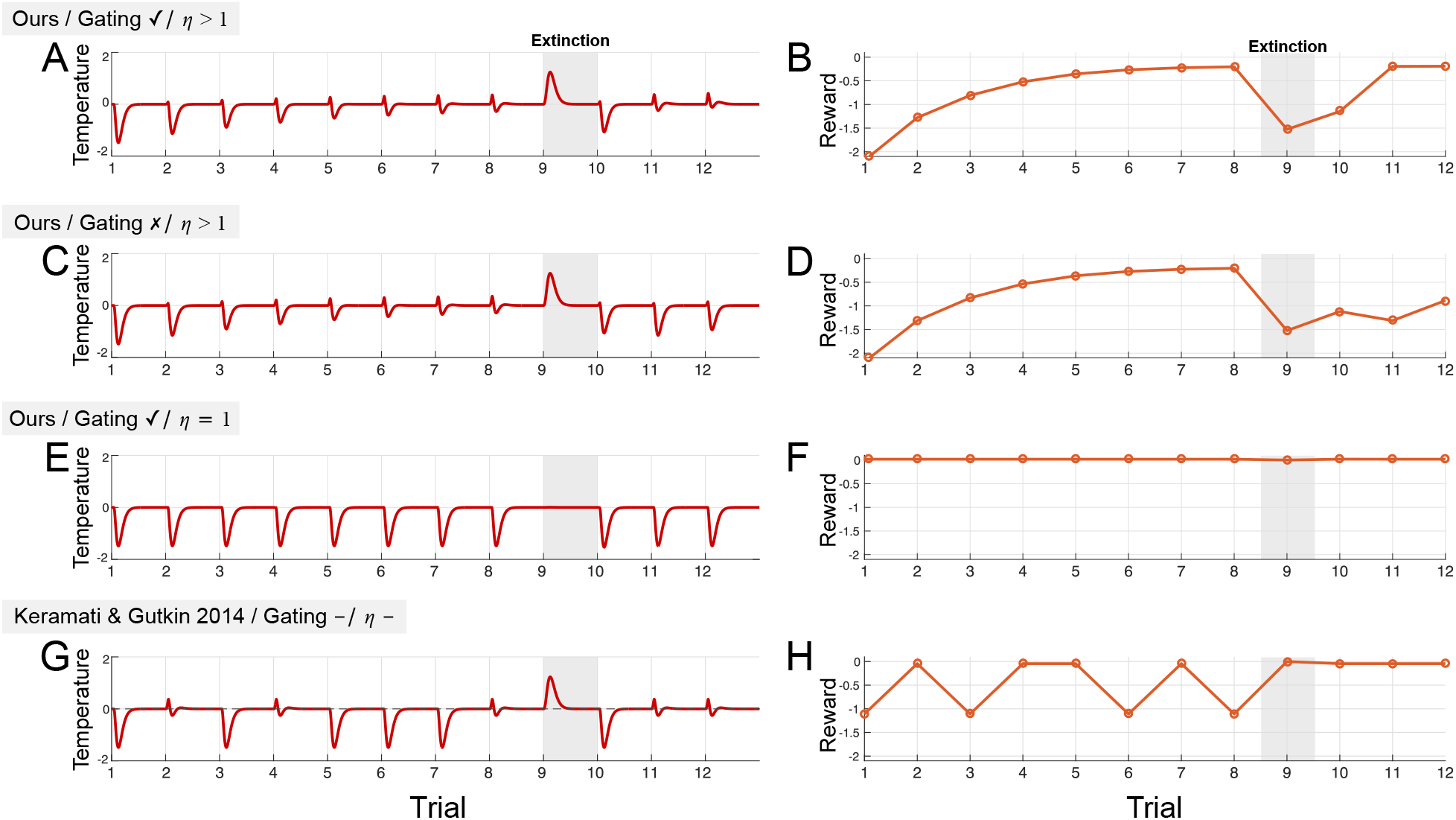
Homeostatic regulation of body temperature. (A, B) Simulated body temperature from normal (A) and net reward (B) produced by the proposed model incorporating a gating function *χ*, which suppresses inhibition during reacquisition trials. The reward weight parameter was set to *η* = 1.5. (C, D) Corresponding results from the same model without the gating mechanism. (E, F) Results with gating intact but with a symmetric reward structure (*η* = 1). (G, H) Results produced by the previous HRL model with discrete all-or-none responses.

Moreover, we checked the behavior after extinction. The proposed model exhibited an immediate recovery of the compensatory response upon reacquisition (Fig. 2A, trial 11), resulting in a sharp increase in reward (Fig. 2B, trial 11). In the lack of the a context-sensitive gating mechanism *χ*, this recovery was absent, even though both activation and inhibition units were present (Fig. 2C and D, trial 11). These results indicate that the gating mechanism plays a critical role in enabling the re-expression of suppressed responses, in addition to the presence of dual learning units. Collectively, our findings support the notion that effective extinction learning requires not only inhibitory memory formation but also a context-dependent mechanism to unmask it.

Next, we examined the role of reward asymmetry. When *η* > 1 (i.e., assigning greater weight to punishment), the model successfully acquired proactive tolerance (Fig. 2A and B). In contrast, when *η* = 1 (i.e., equal weighting of reward and punishment), learning failed (Fig. 2E). This failure arises because the net reward *R* inherently sums to zero (Fig. 2F), as temperature changes that start and return to baseline. These results indicate that imbalanced reward weighting is essential for learning effective regulation.

For comparison, we also implemented the previous model, which allows only binary, all-or-none responses of body temperature in individual trials in a probabilistic manner (Keramati and Gutkin, 2014). This model failed to replicate the gradual development of tolerance (Fig.2G) and showed unstable net rewards across trials (Fig. 2H), highlighting the benefit of our continuous-output model.

### Multi-dimensional model

The one-dimensional (1D) model described above focused on regulating a single variable, namely body temperature, within a framework of HRL. However, real biological systems must maintain homeostasis across multiple variables simultaneously. To capture this complexity, we extended the model to a multi-dimensional framework in which multiple internal variables are regulated via a smaller set of shared meta-signals, such as neural and hormonal activities (Fig. 3A).

**Fig. 3.**
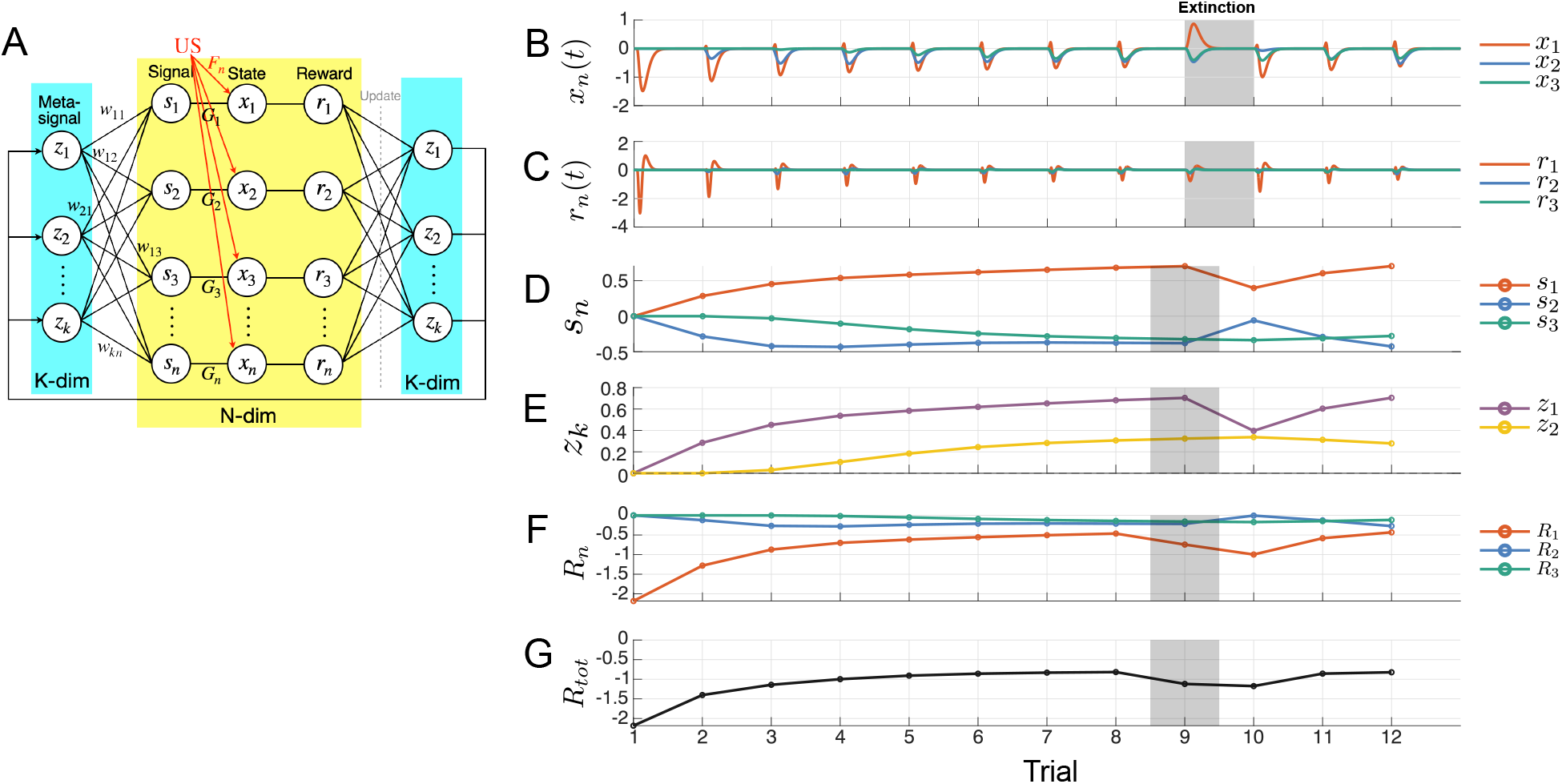
Homeostatic regulation of multiple variables with a meta-signal. (A) Model architecture with *N* local variables and *K* meta-signals. In the simulation, *N* = 3 and *K* = 2. (B-G) Simulated time series of the model variables with the reward weightinf parameter *η* = 1.5. Shaded region indicate extinction phase. (B) The time series of variable *x*_*n*_. The first, second, and third local variables are shown in orange, blue, and green, respectively. (C) Time series of reard *r*_*n*_(*t*). (D) Trial-by-trial changes in local signal strength *s*_*n*_. (E) Meta-signal *z*_1_ (purple) and *z*_2_ (yellow), which modulate *x*_1_, *x*_2_, and *x*_2_, *x*_3_, respectovely. (F) Net rewards *R*_*n*_ for each local variable. (G) Total system reward *R*_tot_.

In biological systems, control signals such as sympathetic tone, vasopressin, or cortisol can simultaneously influence body temperature, blood pressure, hydration, and glucose levels. These shared inputs, referred to here as meta-signals, give rise to trade-offs in regulation: an adjustment intended to stabilize one variable may inadvertently perturb others. In our model, this is captured by allowing each meta-signal to influence multiple variables through fixed regulatory weights *w*.

The extended model consists of *N* internal variables (*x*_1_, *x*_2_, …, *x*_*N*_), each associated with a local control signal *s*_*n*_, which is computed as a weighted sum of *K* meta-signals (*z*_1_, *z*_2_, …, *z*_*K*_) (Fig. 4) as

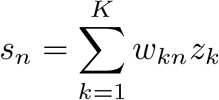

where *N* > *K*, and *w*_*kn*_ is a fixed parameter representing the physiological influence of meta-signal *z*_*k*_ on variable *x*_*n*_. We assume these weights take both positive and negative values, reflecting the fact that meta-signals exert heterogeneous effects. Therefore adjusting one variable toward its optimal value may induce deviations in others. For example, in a hot environment, evaporative cooling through sweating lowers body temperature but leads to water loss. Similarly, pharmacological interventions targeting a specific variable often induce side effects by perturbing others.

**Fig. 4.**
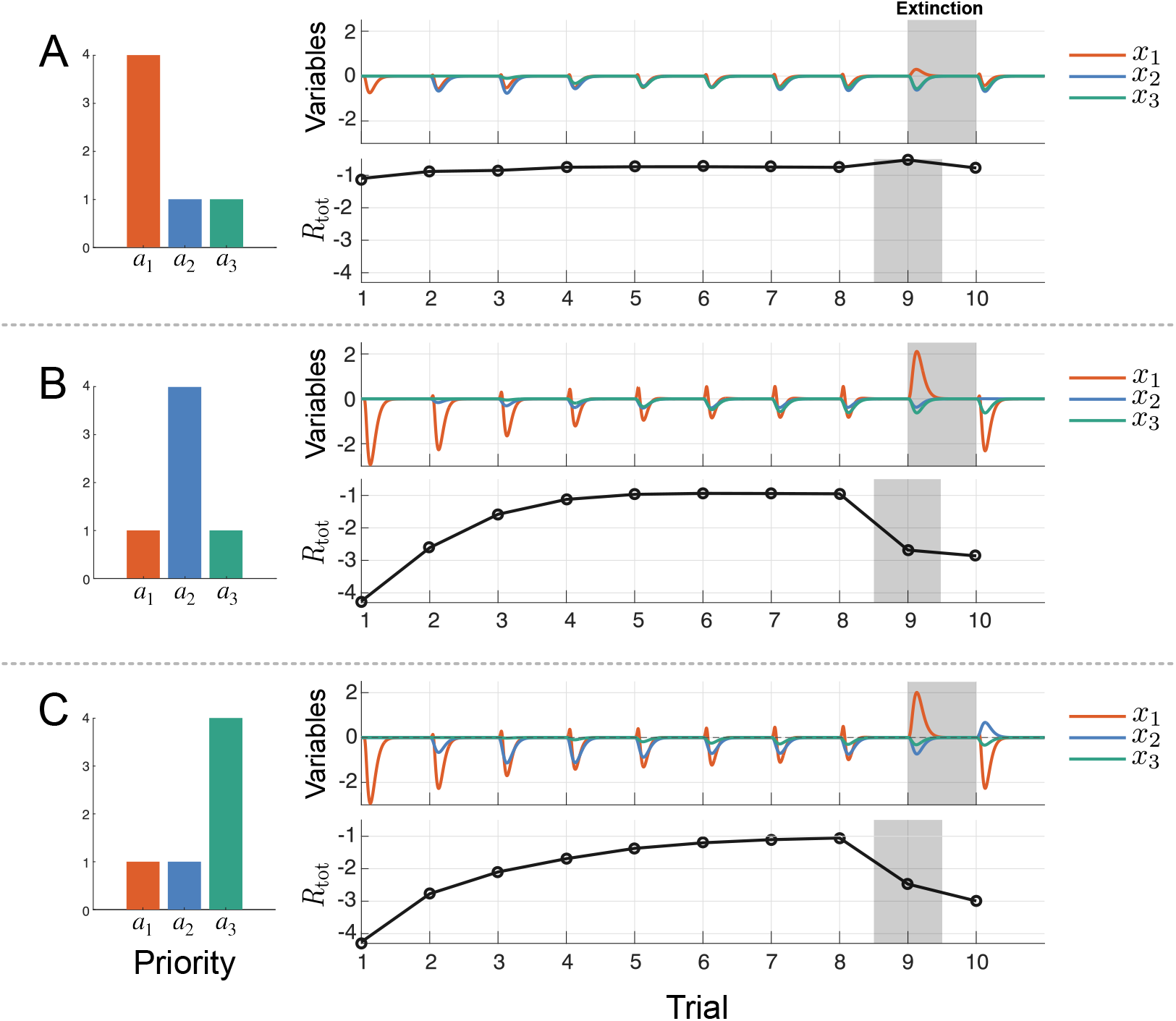
Homeostatic regulation of multiple variables under three priority weight patterns. (A–C) Simulations with three different priority configurations, where variable *x*_1_ (A), variable *x*_2_ (B), or variable *x*_3_ (C) is assigned the highest weight, as shown in the bar graphs (left). Red, green, and blue bars represent priority weights for variables *x*_1_, *x*_2_, and *x*_3_, respectively. For each configuration, the top-right panel shows the time series of deviations from each setpoint *x*_*n*_(*t*) − 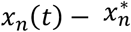 and the bottom-right panel shows the corresponding total reward *R* across trials.

The dynamics of each internal variable *x*_*n*_ are described by

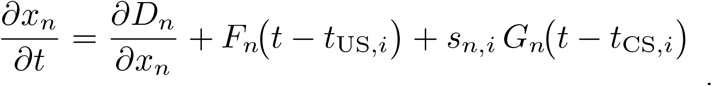

The drive function, defined as 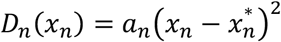, quantifies the motivational force to restore *x*_*n*_ to its fixed setpoint 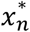, where an represents priority assigned to each variable. *F*_*n*_ (*t*) and *G*_*n*_ (*t*) denote the response profiles to US and CS, respectively. *t*_US,*i*_ and *t*_CS,*i*_ are the timings of unconditioned and conditioned stimuli in the *i*-th trial.

Learning in this multi-dimensional HRL model proceeds by updating the meta-signals *z*_*k*_ to maximize the cumulative reward of the local variables they influence. Since each meta-signal affects multiple variables, and each variable has its own priority weight *a*_*n*_, the reward function implicitly integrates across all variables under shared control. As a result, optimization of meta-signals involves navigating trade-offs among competing variables, favoring adjustments in higher-priority variables in order to improve the overall reward.

### Simulation of Proactive Regulation of the Multiple States

To explore the dynamics of the multidimensional model and its capacity to maintain homeostasis across multiple variables, we focused on two key factors influencing the system’s behavior: (1) prioritization among local states and (2) degree of imbalance weight on positive and negative reward. To examine these effects, we considered the model comprising three variables *x*_1_, *x*_2_, *x*_3_(*N* = 3), where *x*_1_ was designated as the target of the external disturbance (US). The model included two meta-signals *z*_1_ and *z*_2_(*K* = 2). The physiological influence matrix of meta-signal, *w*, was given as

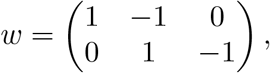

which represents their up- and down-regulatory effects on the local variables in the multidimensional simulation. Although each variable *x*_*n*_ in the model is an abstract representation of an internal physiological state, this formulation can be interpreted as a simplified representation of competing homeostatic dimensions such as body temperature, plasma glucose level, and hydration state, which are known to interact through shared regulatory signals.

#### (1) Prioritization among local states

Each variable xn represents a distinct physiological state with its own optimal setpoint 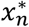. The prioritization for each variable is encoded in the priority parameter an within the drive function. Depending on *a*_*n*_, some variables are vital for survival and thus tightly maintained near their setpoints, while others can tolerate temporary deviations without substantially disrupting overall homeostasis. We examined model behavior under two prioritization patterns.

We simulated the case with uniform prioritization pattern (*a*_*n*_ = 2 for all *n*) (Fig. 3B-G). The system gradually improved x over time (Fig. 3B). Although external disturbances were applied only to *x*_1_(*t*), *x*_2_(*t*) deviated from its optimal value 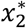 due to the influence of the meta-signal *z*_1_, which jointly regulates *x*_1_ and *x*_2_. Subsequently, *x*_3_ deviated from its own optimum to compensate for the accumulated deviations. Thus, local variables adjusted their setpoints to maximize the system-wide global reward *R*_tot_ (Fig. 3G).

We also simulated the model with non-uniform prioritization configurations, in which each of the three variables was assigned the highest priority in turn. When *x*_1_, the target of external disturbance, held the highest priority (i.e., *a*_1_ > max(*a*_2_, *a*_3_)), the system exhibited only minor perturbation due to stronger reactive regulation mediated by the drive term *∂D*_1_/*∂x*_1_. As a result, the disturbed state rapidly returned to its setpoint (Fig. 4A). In this condition, the deviation of *x*_1_ was effectively minimized, although deviations in *x*_2_ and *x*_3_ emerged as compensatory trade-offs. Consequently, the global reward *R*_tot_ increased only modestly. In contrast, when either *x*_2_ or *x*_3_ was prioritized (i.e., *a*_2_ > max(*a*_1_, *a*_3_) or *a*_3_ > max(*a*_1_, *a*_2_)), the initial perturbation in *x*_1_ was larger and its recovery was slower. This resulted in delayed reward accumulation and greater compensatory deviations in the other variables (Fig. 4B, C). Overall, fluctuations in the higher-priority variables were effectively suppressed, while disturbances affecting lower-priority variables were tolerated and left largely uncorrected. Accordingly, when either *x*_2_ or *x*_3_ received the highest priority, the system failed to fully compensate for the the external disturbances applied to *x*_1_, reflecting a prioritization strategy that protects critical variables at the expense of others.

#### (2) Joint parameter effect on RL success

We systematically investigated how reward asymmetry (*η*) and prioritization (*a*) jointly influence the learning dynamics. The prioritization pattern was parameterized by a single scalar angle, such that 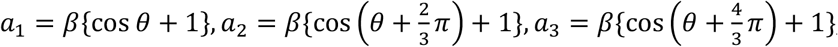, which allows continuous variation across the entire spectrum of prioritization configurations (Fig.5A). For each parameter pair (*η, θ* ), we performed a grid-search to identify the optimal strength of the meta-signals ***z***^*^(*η, θ*), ***z*** = (*z*_1_, *z*_2_, …, *z*_*K*_), that maximizes the system’s reward in a trial, *R*_tot_ (Fig. 5B, Fig. S2).

**Fig. 5.**
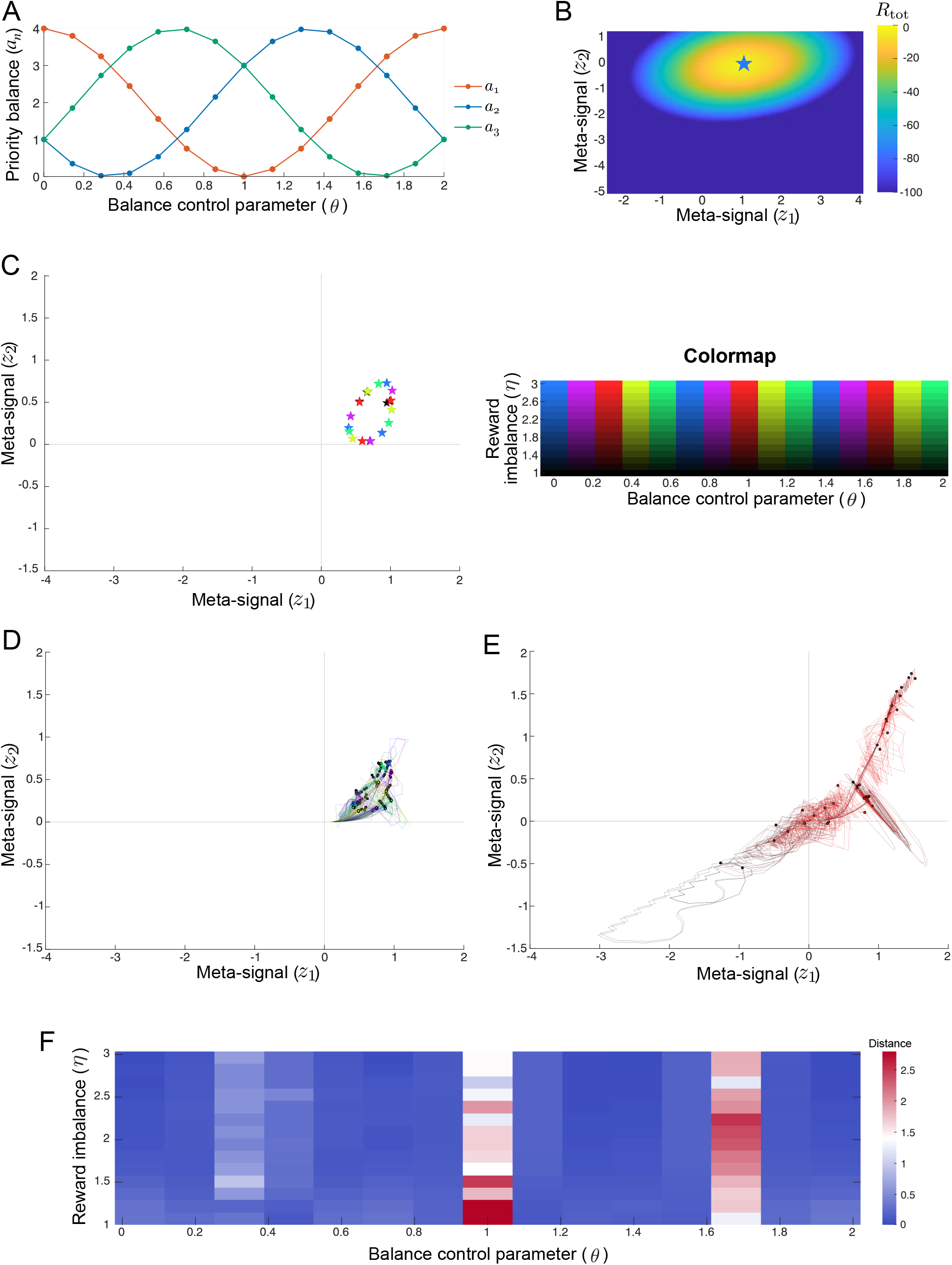
Influence of parameter pairs on multidimensional model dynamics. (A) Parameterization of priority weights *a*_*n*_ as a function of prioritization angle *θ*. Orange, green, and blue lines represent *a*_1_, *a*_2_, *a*_3_, respectively. (B) Example heatmap of *R*_tot_ depending on meta-signal strengths. The star represents the reward optimal meta-signal ***z***^*^(*η, θ*), where *η* = 3 and *θ* = 1.3878. (C) Reward-optimal ***z***^*^(*η, θ*) for each parameter pair. Each star marks the reward optima ***z***^*^(*η, θ*); hue denotes *θ* and lightness denotes *η* (see the right panel). As the optimum’s location varies mainly with *θ*, points for lower *η* (darker shades of the same hue) lie directly behind those for higher *η* and may not be visible. (D, E) Trajectories of the two meta-signals (*z*_1_, *z*_2_) across 100 trials for representative values of *θ*. Trajectories were smoothed by averaging across every 3 trials in (D) balanced prioritization conditions (*θ* ∉ {0.2857, 0.9796, 1.6735}) and (E) disproportionate prioritization conditions(*θ* ∈ {0.2857, 0.9796, 1.6735}). (F) Heatmap of the mean distance to the reward optimum ***z***^*^(*η, θ*) across the (*η, θ*)-plane (averaged over the final 10 trials).

Because RL success or failure can be evaluated by whether the simulated meta-signal trajectory ***z***_*k,i*_ approaches the reward-optimal ***z***^*^(*η, θ*), we simulated the multidimensional model for 100 CS–US trials without an external disturbance. This procedure resolved the trajectory of the meta-signals under each parameter setting (Fig. 5D). When the prioritization was more balanced, trajectories remained tightly confined around the optimum (Fig. 5D left); by contrast, under extremely disproportionate prioritization, trajectories deviated from the optimum (Fig. 5D right).

Lastly, we tested whether the meta-signals reliably converge to the reward optimum ***z***^*^(*η, θ*) across the (*η, θ*) space. Deviation from the optimum was quantified as the mean Euclidean distance between the simulated trajectory and ***z***^*^(*η, θ*) over the final 10 trials. This systematic sweep revealed contiguous regions of the (*η, θ*)plane where the meta-signals consistently converged to ***z***^*^(*η, θ*), and distinct regions where trajectories moved away from the optimum, indicating failure to learn appropriate compensatory responses (Fig. 5E). The resulting phase diagram delineates the parameter regimes in which the proposed RL mechanism is expected to succeed. Importantly, the external disturbance acted directly only on the first variable, *x*_1_ (Fig. 3A)

## Discussion

### Summary

This study proposes an extended framework of homeostatic reinforcement learning (HRL) that enables proactive and gradual regulation of internal physiological states through experience. In a one-dimensional thermoregulatory conditioning task, the model reproduces ethanol tolerance and rapid reacquisition after extinction by learning trial-by-trial compensatory responses, consistent with CS–US thermoregulatory data (Mansfield and Cunningham, 1980) Extending to multiple physiological axes, the framework shows that shared regulatory signals impose unavoidable trade-offs: priority structure determines which variables are tightly stabilized and which are allowed to fluctuate, thereby shaping convergence or failure.

### Extensions beyond the original HRL Model

Relative to the original HRL model (Keramati and Gutkin, 2014), three biologically informed ingredients are central and each is needed to capture the observed phenomena. First, continuous control signals allow graded, trial-by-trial compensation rather than all-or-none actions. In the one-dimensional thermoregulatory task, this continuity reproduces gradual tolerance and rapid reacquisition after extinction, in line with CS–US thermoregulatory findings and Pavlovian tolerance reports (Farahbakhsh and Siciliano, 2023; Mansfield and Cunningham, 1980; Poulos and Cappell, 1991; Siegel, 2005,2001). Second, loss-dominant valuation – greater weight on increases in drive than on equivalent decreases – is required in our formulation. Without this asymmetry, the temporal-derivative reward becomes uninformative and prevents learning. This is consistent with asymmetry observed in biological decision-making (Macpherson et al., 2014; Sokol-Hessner and Rutledge, 2019). Third, inhibitory learning with context-sensitive gating treats extinction as inhibition of expression rather than of the original memory trace. This representation aligns with evidence that extinction recruits inhibitory circuits and is context dependent, which in turn allows re-expression phenomena such as spontaneous recovery and rapid reacquisition (Bouton, 2002; Bouton et al., 2006; Quirk and Mueller, 2008; Whittle et al., 2021).

In the multidimensional extension, the framework jointly regulates several variables through shared meta-signals and explicitly specified priority weights. This arrangement clarifies how priority choices create trade-offs across axes. Prioritizing one variable tightens stabilization of that axis at the expense of others.

### Pathological Relevance

In the multidimensional model, when one homeostatic variable receives disproportionately low priority, its deviation is effectively ignored during learning, leading to persistent misattribution of disturbances as externally caused and to non-convergence of compensatory responses. Over time, such model dynamics suggest a cautious link to autonomic dysregulation. In myalgic encephalomyelitis/chronic fatigue syndrome (ME/CFS), for example, orthostatic intolerance and related autonomic abnormalities are frequently reported (Freeman and Komaroff, 1997; Garner and Baraniuk, 2019) and reduced heart-rate variability (HRV) indicates altered cardiac autonomic regulation (Nelson et al., 2019), with HRV abnormalities further associated with fatigue severity (Escorihuela et al., 2020). These empirical features are consistent with theoretical accounts emphasizing impaired allostatic control and aberrant interoceptive inference (Barrett and Simmons, 2015; Stephan et al., 2016). In this framing, clinical manifestations can be mapped onto specific parameter regimes of the model, thereby linking clinical observations to the computational dynamics of HRL. Perturbation-learning protocols could be used to estimate relative priority weights from time-series data and to assess the prevalence of non-convergent adaptation. Such tests would evaluate the model’s relevance to patient populations.

### Comparison with Prior Works

Classical homeostatic models grounded in negative feedback have been highly successful at explaining rapid, reactive stabilization of internal variables—for example, baroreflex control of blood pressure and thermoregulatory set-point maintenance (Ashby, 2013; Cannon, 1932). By design, however, these formulations do not incorporate experience-dependent learning or contextual adaptation, and thus do not readily account for gradual tolerance or cue-dependent modulation of internal states.

A complementary perspective is allostasis, which emphasizes “stability through change,” highlighting predictive, context-sensitive adjustments of set points (Sterling & Eyer 1988)(McEwen, 2017; Ramsay and Woods, 2014; Sterling, 2012, 2004). This literature provides conceptual and empirical motivation for anticipatory regulation in stress and neuroendocrine systems, but it has largely remained descriptive and does not specify how such predictive adjustments are learned from repeated experience.

Predictive coding and the free-energy principle offer a normative route to anticipatory control, casting agents as inference systems that maintain and update generative models of body and environment to minimize prediction error (Friston et al., 2015; Friston and Stephan, 2007; Pezzulo et al., 2024; Tschantz et al., 2022). These approaches have broad explanatory scope but typically assume structured priors or learned latent dynamics and entail nontrivial computational demands, which may be challenging for fast, resource-limited autonomic regulation.

Our reinforcement-learning account complements these traditions by producing anticipatory regulation without constructing explicit generative models. Building on prior HRL work (Keramati and Gutkin, 2014), we introduce biologically informed components that together reproduce gradual tolerance, extinction with re-expression (spontaneous recovery/rapid reacquisition), and multivariable trade-offs. In this way, the framework provides a learning-based, model-free complement to predictive coding/FEP and a computational instantiation of allostatic ideas, while remaining simple enough to yield testable predictions about when regulation converges or fails (Friston et al., 2015; Friston and Stephan, 2007; Ramsay and Woods, 2014; Sterling, 2012).

### Limitations and Future Directions

Despite its strengths, the model has limitations. Reward was defined solely as the temporal derivative of the drive function, reducing sensitivity to persistent deviations. The influence matrix *w* was fixed, limiting flexibility in representing adaptive reweighting of regulatory signals. Moreover, simulations were not fit directly to multi-variable physiological datasets. Future work should integrate empirical data (e.g., temperature, glucose, hydration) and allow adaptive coupling weights to capture long-term physiological changes. Parameter regimes that yielded instability in simulations may correspond to dysregulated states in real organisms; testing these predictions against pathological data represents an important next step.

Beyond these technical considerations, the framework offers a compact yet extensible bridge between reinforcement learning and physiological regulation. By emphasizing experience-dependent adaptation within biologically realistic constraints, it complements control-theoretic, Bayesian, and predictive coding approaches. Looking forward, its integration with empirical datasets may transform it from a conceptual model into a translational tool—deepening our understanding of adaptive and maladaptive regulation, and guiding strategies for intervention.

## Methods

### 1D model

The dynamics of body temperature x in the i-th trial are described by the equation:

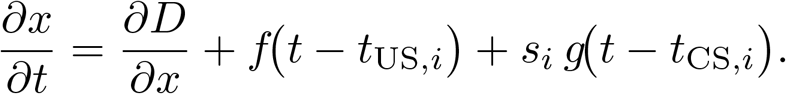

*f*(*t*) and *g*(*t*) are transient functions which start at *t* = 0, where *f*(*t*) = 0 and *g*(*t*) = 0 when *t* ≤ 0. *t*_US_ and *t*_CS_ denote the timings of US and CS, respectively. The function *f*(*t*) represents the effect of ethanol injection on body temperature, leading to hypothermia, while *g*(*t*) represents the compensatory response. In our simulation, the dynamics of body temperature were modeled using this equation for each trial, which was repeated multiple times.

The drive function *D*+*x*(*t*)) represents the motivation to shift the body state based on the current state, which is defined as the distance of the internal state from the desired setpoint *x*^*^. In this study, we employ a quadratic function

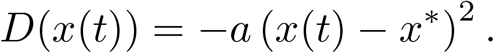

The model optimizes the control variable sn to maximize reward r. In the context of homeostatic reinforcement learning, the reward at each time t is defined as the negative temporal derivative of the drive function D, i.e., dD/dt=a(x-x*)dx/dt. Additionally, we introduced an asymmetry in positive and negative rewards based on prospect theory, by adding a weight on negative reward (punishment). The instant reward r(t) is defined as

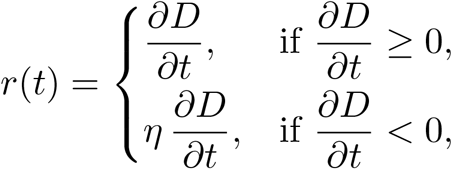

where >1, which represents the asymmetry. The net reward of the *i*-th trial *R*_*i*_ and the cumulative deviation of body temperature *X*_*i*_ were calculated as temporal integrals of their time serieses

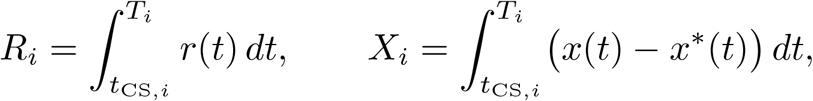

where *T*_*i*_ denotes the time that the *i*-th trial ends. Note that *R*_*i*_ is always negative and zero if *η* > 1 and *η* = 1, respectively. The strength of the compensatory signal *s*_*i*_ is assumed to be modulated by two neural populations, *A* and *I*, which respectively promote and suppress the response both driven by the CS. Their interaction is governed by a gating variable *χ*_*i*_ (i.e., *χ*_*i*_ = CS_i-1_US_i-1_, CS_i_,US_i_ ∈ {0,1}), defined as

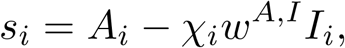

where *w*^*A,I*^ represents the inhibitory effect of *I* on *A*, and indicates the effect size of neural activity on temperature dynamics. This gating mechanism reflects the idea that an activatory memory can be masked by an inhibitory trace, and that such inhibition can be transiently desengaged upon presentation of a retrieval cue. This formulation is consistent with the biological observation that extinction does not erase original associative learning, but instead superimposes context-dependent inhibition. Both *A* and *I* are upregulated by CS input

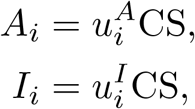

where 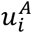 and 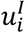 are weights on *A*_*i*_ and *I*_*i*_, respectively. These weights are updated based on the discrepancy between actual and ideal rewards, following the assumptions introduced in the Rescorla-Wagner model, a classical conditioning model. The update of signal weights 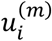 is

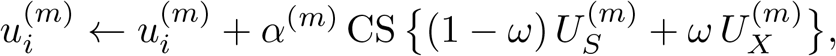

where *α* represents the learning rate and *m* ∈ {*A, I*}.

The update process is divided into two components: 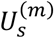 and 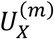, which represent how much reward is predicted by self-regulation of 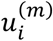 and by unexpected deviations of the body state e.g., when the acquisition phase changes to the extinction phase and vice versa. 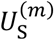 is described by temporal reward shift *R*_*i*_ with respect to the increment of 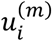, with a sigmoid function *h* to limit the update amount per step, as

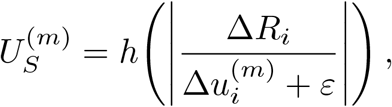

where 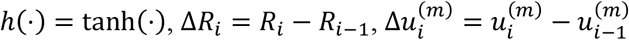 and *ϵ* is the small constant to avoid zero division. In contrast, 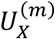 is described by temporal change in reward *R*_*i*_ with respect to the deviation of *X*_*i*_, as

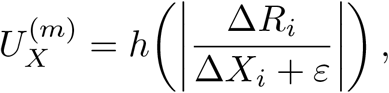

WhereΔ*X*_i_= *X*_i_ − *X*_*i*− *1*_

In the second term, *ω* (0 ≤ *ω* ≤ 1) determines the weight of 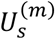 and 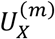 as

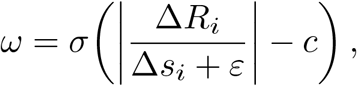

where *σ*(·) = 1/(1 + exp(·)), Δ*s*_*i*_ = *s*_*i*_ − *s*_*i*–1_, and *c* denotes constant parameter. This *ω* implies how likely the cause of the change in *R*_*i*_ was either a self-regulation signal or unexpected deviations of the body state (For intuition, we visualize the functional form of *σ* relative to an example dataset: see Supplementary Fig. S1). When *R*_*i*_ improves gradually as a result of the update of signal weights 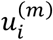, takes a small value hence 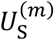 is dominant. In contrast, when *R*_*i*_ experiencing sudden changes (jump or drop) due to unexpected deviations of the body state, *ω* takes a large value hence 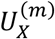 is applied. In this scenario with large *ω*, since the change in *R*_*i*_ is not directly related to 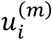, the gradient for updating 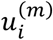 is approximated by calculating the temporal difference of net reward *R*_*i*_ with respect to the deviation of *X*_*i*_.

### Multi-dimensional extension

In the one-dimensional (1D) model described above, the regulation of a single internal variable, body temperature, was considered. However, real biological systems must simultaneously maintain homeostasis across multiple physiological variables.

Here, we consider a scenario in which the brain controls multiple internal variables (e.g., body temperature, blood pressure, blood glucose level) through a limited number of command signals, such as peripheral neural activity or hormonal secretion. In such a case, each command signal influences multiple variables simultaneously. Consequently, adjusting one variable toward its optimal value may cause other variables to deviate from theirs. For instance, when the body lowers an elevated temperature via sweating, the body temperature approaches its optimal range, but this leads to a loss of water. Likewise, pharmacological interventions often have side effects: while a drug may alleviate one condition, it may simultaneously disturb another.

Within the framework of Homeostatic Reinforcement Learning (HRL), where each internal variable is associated with a fixed set point, a single action that optimizes one variable does not necessarily bring all other variables closer to their respective set points. Regulatory actions therefore involve inevitable trade-offs, producing both beneficial and adverse consequences.

At each (*i*-th) trial, each variable *x*_*n,i*_ is regulated by a local signal *s*_*n,i*_ which is in turn influenced by a weighted sum of meta-signal *z*_*k,i*_

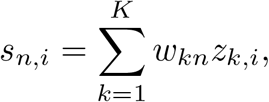

where *w*_*kn*_ represents the fixed physiological weight from the *k*-th meta-signal to the *n*-th variable (see Fig. 3)

The dynamics of the *n*-th variable *x*_*n*_ are defined as

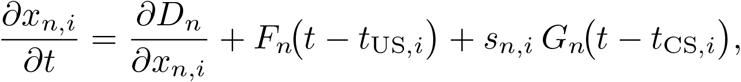

where is the drive function, *t*_US,*i*_ and *t*_CS,*i*_ denote the timings of conditional (CS) and unconditioned stimuli (US), respectively, and *F*_*n*_(*t*) and *G*_*n*_(*t*) are US- and CS-induced input functions for the n-th variable. The amplitude of each *G*_*n*_(*t*) is modulated by the local signal *s*_*n,i*_ which itself is regulated by the meta-signals *z*_*k,i*_.

An instantaneous reward *r*_*n*_(*t*) is defined independently for each variable. As in the 1D model, the net reward *R*_*n,i*_ and the cumulative deviation of variable *X*_*n,i*_ for the *i*-th trial are computed accordingly as follows.

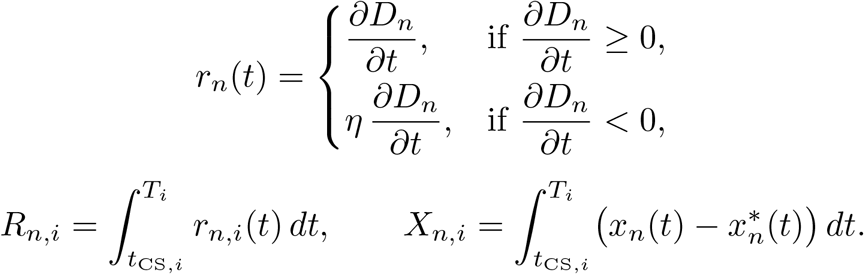

The meta-signal *z*_*k*_ is optimized through a control variable *u*_*k*_, which aims to maximize the total reward 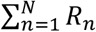. The meta-signal *z*_*k*_ is assumed to be modulated by two neural populations, *A*_*k*_ (excitatory) and *I*_*k*_ (inhibitory), with the control variable 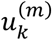 (for *m* ∈ {*A, I*}) updated as

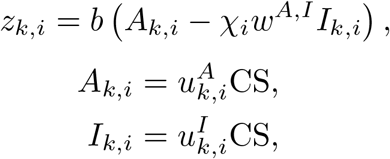

with a gating variable *χ*_*i*_ (i.e., *χ*_*i*_ = CS_i-1_US_i-1_, CS_i_,US_i_ ∈ {0,1}). The parameter *w*^*A,I*^ represents the inhibitory effect of *I*_*k*_ on *A*_*k*_, and indicates the effect size of neural activity on temperature dynamics.

The update consists of two components: 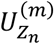, which reflects self-regulation based on predicted reward changes, and 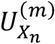, which reflects adjustments due to external disturbances. These are defined as

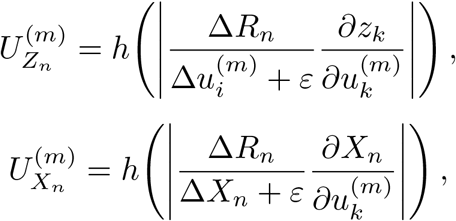

where 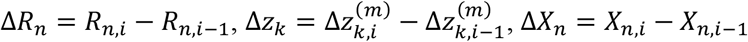 and *ϵ* is a small constant to avoid division by zero.

The weighting parameter *ω*_*n*_, which determines the relative contribution of 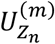 to 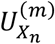, is computed as

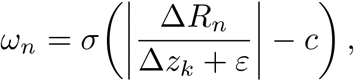

where *c* is a parameter that shifts the sigmoid function. All model parameters are listed in Table 1 in Supprementary.

## Acknowledgements

We are grateful to Prof. Michiyuki Matsuda and Prof. Kazuhiro Aoki for providing research environment. We thank Sana Ishihara for assistance with preliminary model construction related to this study.

This study was supported in part by Japan Society for the Promotion of Science (JSPS) KAKENHI [23KJ1293 to M.F], Japan Science and Technology Agency (JST) the Moonshot R&D–MILLENNIA

Program [JPMJMS2024-9 to H.N.], and Japan Agency for Medical Research and Development (AMED) Multidisciplinary Frontier Brain and Neuroscience Discoveries (Brain/MINDS 2.0) [JP25wm0625322 to H.N.].

## Author contributions

M.F. and H.N. conceived the research project and wrote a manuscript. M.F. developed the model and implemented analysis.

