## Supplemental Fig. S1, Fig. S2, Table 1. for "Gradual proactive regulation of body state by reinforcement learning of homeostasis"

### Supplementary

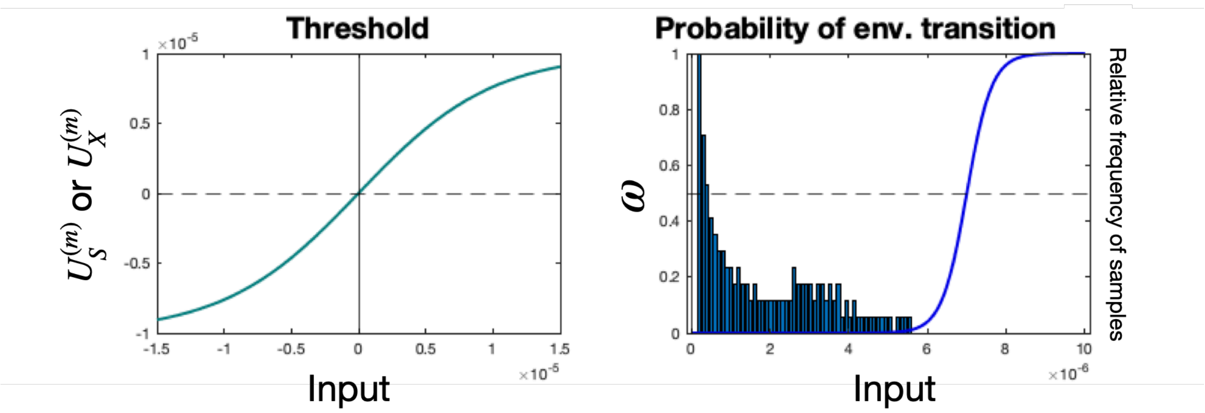


Fig. S1. Visualization of $h(x) = C_1 \tanh(C_2 x)$ and $\sigma(x) = 0.5 + C_3 \tanh(C_4 x)$ relative to a representative dataset. (A) The thresholding function $h$, computing $U_S^{(m)}$ and $U_X^{(m)}$. (B) The sigmoid function $\sigma$ (blue line), which computes the probability of environmental transition $\omega$, and a histogram of sample observations fitted to the scale of $\sigma$. Axes and parameter settings match those used in the main analyses.

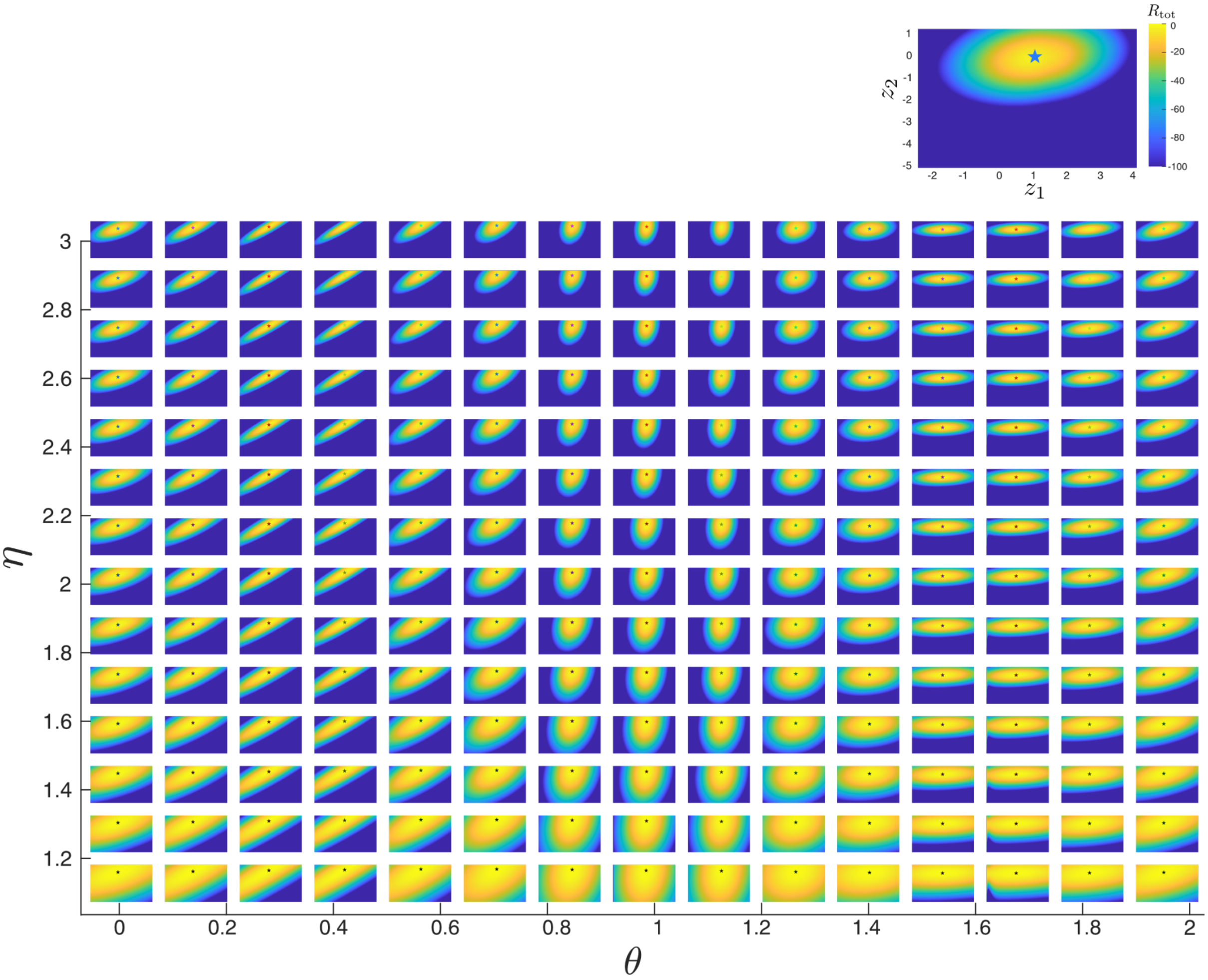


Fig. S2. **Heatmaps of $R_{\text{tot}}$ across meta-signal strengths for each parameter pair.** For each combination of reward asymmetry ($\eta$) and prioritization angle ($\theta$), $R_{\text{tot}}$ was evaluated over a two-dimensional grid of meta-signal strengths ($z_1, z_2$). We sampled 15 values of uniformly over [0, 2) and 15 values of uniformly over [1,3], yielding a 15 x 15 grid (225 settings), and 65 samples of $z_1$ and $z_2$ (4425 samples) for grid-search, using a learning rate ($A = 1.5, I = 4$; fixed across runs). The star in each panel indicates the location of the reward-optimal meta-signal. The colour of star corresponds to $\eta$-$\theta$ parameter pair colour map in Fig. 5C.

Table 1. Parameter settings

| Parameter | Value |
|---|---|
| $\alpha^A$ | 1.5 |
| $\alpha^I$ | 4 |
| $b$ | 2 |
| $w^{A,I}$ | 1 |
| $\beta$ | 2 |
| $c$ | -0.03 |
| $C_1$ | 0.1 |
| $C_2$ | 0.1 |
| $C_3$ | 0.5 |
| $C_4$ | 30 |
